# Sulforaphane exhibits in vitro and in vivo antiviral activity against pandemic SARS-CoV-2 and seasonal HCoV-OC43 coronaviruses

**DOI:** 10.1101/2021.03.25.437060

**Authors:** Alvaro A. Ordonez, C. Korin Bullen, Andres F. Villabona-Rueda, Elizabeth A. Thompson, Mitchell L. Turner, Stephanie L. Davis, Oliver Komm, Jonathan D. Powell, Franco R. D’Alessio, Robert H. Yolken, Sanjay K. Jain, Lorraine Jones-Brando

## Abstract

Severe Acute Respiratory Syndrome Coronavirus 2 (SARS-CoV-2), the cause of coronavirus disease 2019 (COVID-19), has incited a global health crisis. Currently, there are no orally available medications for prophylaxis for those exposed to SARS-CoV-2 and limited therapeutic options for those who develop COVID-19. We evaluated the antiviral activity of sulforaphane (SFN), a naturally occurring, orally available, well-tolerated, nutritional supplement present in high concentrations in cruciferous vegetables with limited side effects. SFN inhibited in vitro replication of four strains of SARS-CoV-2 as well as that of the seasonal coronavirus HCoV-OC43. Further, SFN and remdesivir interacted synergistically to inhibit coronavirus infection in vitro. Prophylactic administration of SFN to K18-hACE2 mice prior to intranasal SARS-CoV-2 infection significantly decreased the viral load in the lungs and upper respiratory tract and reduced lung injury and pulmonary pathology compared to untreated infected mice. SFN treatment diminished immune cell activation in the lungs, including significantly lower recruitment of myeloid cells and a reduction in T cell activation and cytokine production. Our results suggest that SFN is a promising treatment for prevention of coronavirus infection or treatment of early disease.

## Introduction

The multi-functional phytochemical sulforaphane (SFN) is the isothiocyanate derived from enzymatic hydrolysis of its precursor glucoraphanin, a glucosinolate found in high concentrations in broccoli (*Brassica oleracea italica*). SFN is a potent naturally occurring activator of the transcription factor nuclear factor erythroid 2-related factor 2 (NRF2), with well-documented antioxidant and anti-inflammatory effects ^1,2^. Treatment with SFN increases phagocytic activity of alveolar macrophages ^3^ and reduces lung injury in animal models of acute respiratory distress syndrome (ARDS) ^4^. SFN also lowers the levels of IL-6 and viral load in human subjects infected with live attenuated influenza virus ^5,6^. Numerous clinical trials utilizing SFN demonstrate favorable pharmacokinetics after oral dosing and document excellent tolerability and safety ^1,7,8^.

Our aim was to interrogate SFN for efficacy against human coronaviruses. We report here that SFN inhibits in vitro seasonal coronavirus HCoV-OC43 and Severe Acute Respiratory Syndrome Coronavirus 2 (SARS-CoV-2) infections of mammalian host cells and appears to have a synergistic interaction with remdesivir. In addition, SFN reduces viral load and pulmonary pathology in a mouse model of SARS-CoV-2 infection.

## Results

To evaluate the potential virus-inhibitory activity of SFN, cells were exposed in vitro to the test drug for 1 – 3 hours before inoculation with coronaviruses. In this near-simultaneous drug-infection scenario, SFN effectively inhibited both HCoV-OC43 and SARS-CoV-2-Wuhan-Hu-1 virus-associated cell death in non-human primate Vero C1008 cells in a dose-dependent manner revealing comparable median inhibitory concentrations (IC_50_ = 10 μM, 95% CI 4.7 – 20.4, and 12 μM, 95% CI 4.7 - 30, respectively), and virus selectivity (TI_50_ = 7 and 7, respectively) (Figures 1A, 2A). When the same assay was performed using human diploid fibroblasts, MRC-5, SFN treatment of HCoV-OC43 infection produced similar results (IC_50_ = 18 μM, 95% CI 9.7 −33.5, TI_50_ = 5) (Figure 1B). SFN cytotoxicity was also dose-dependent, the median cytotoxic dose (TD_50_) remained within the range of 73 – 89 μM (Figures 1A-B, 2A). SARS-CoV-2-associated cytopathogenicity was not evaluated in human cells because viral infection did not result in measurable cell death. Instead, we quantified viral RNA from SARS-CoV-2 infected human intestinal Caco-2 cells treated with SFN. A dose-dependent reduction was observed with an IC_50_ of 2.4 μM (Figure 2C).

**Figure 1.**
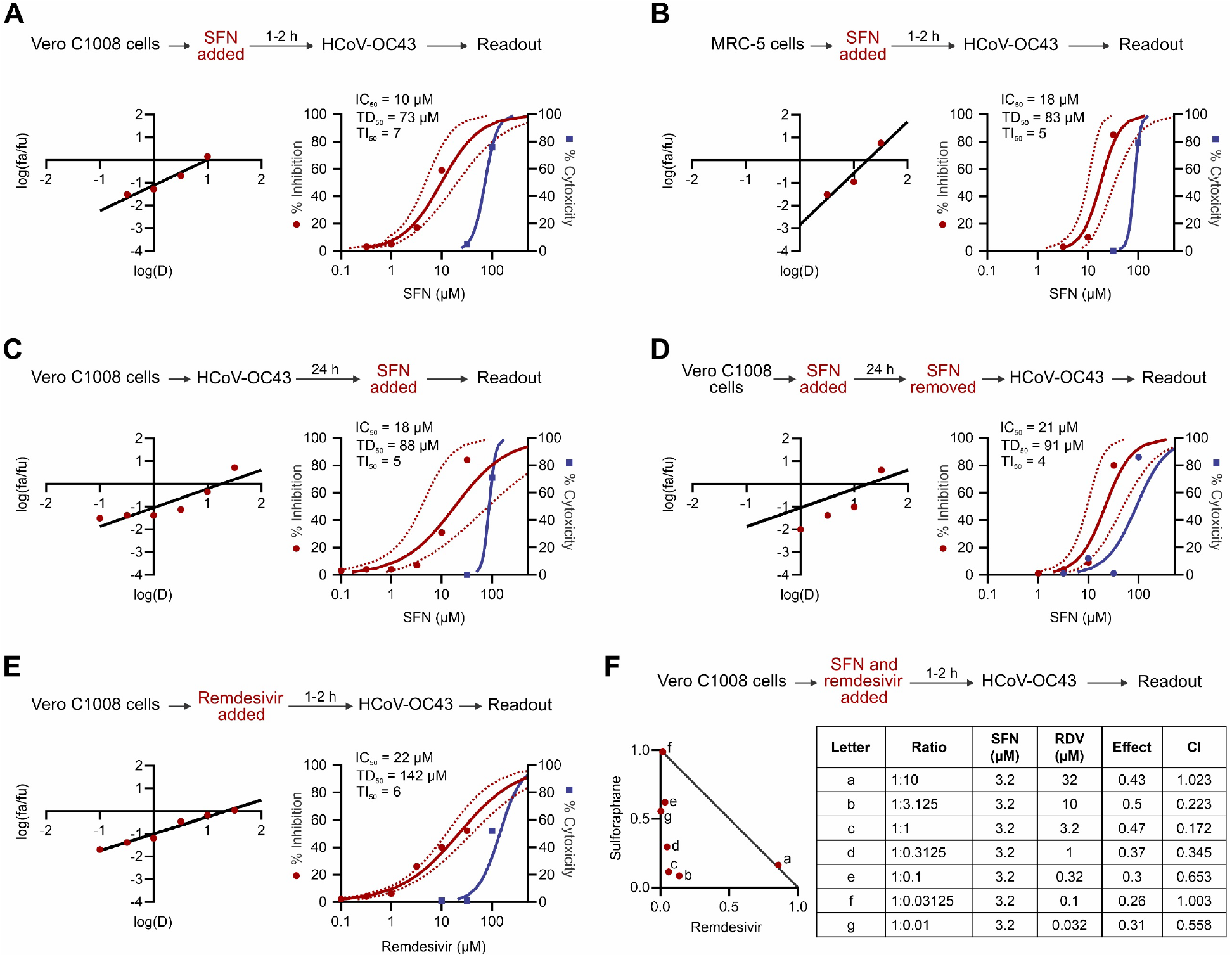
Antiviral effects of SFN against HCoV-OC43. Median effect plots and dose-effect curves calculated for (**A**) Vero C1008 cells infected with HCoV-OC43 after a 1-2 h incubation with increasing concentrations of SFN; (**B**) MRC-5 cells infected with HCoV-OC43 after a 1-2h incubation with increasing concentrations of SFN; (**C**) Vero C1008 cells infected with HCoV-OC43 over 24 h, after which they were incubated with SFN; (**D**) Vero C1008 cells incubated with SFN for 24 h, after which the drug was removed and the cells were infected with HCoV-OC43; (**E**) Vero C1008 cells infected with HCoV-OC43 after a 1-2h incubation with increasing concentrations of remdesivir. (**F**) Normalized isobologram showing combination index (CI) for combinations of various doses. Antiviral data displayed in red; anti-host cell activity (cytotoxicity) displayed in blue. Synergism (CI < 1); Additive effect (CI=1); Antagonism (CI >1); SFN, Sulforaphane; RDV, remdesivir. Dotted lines represent 95% confidence interval. Experiments performed a minimum of 2 times (range = 2 – 7), 3 – 6 replicates within each experiment, except experiment shown in D which was performed once.

**Figure 2.**
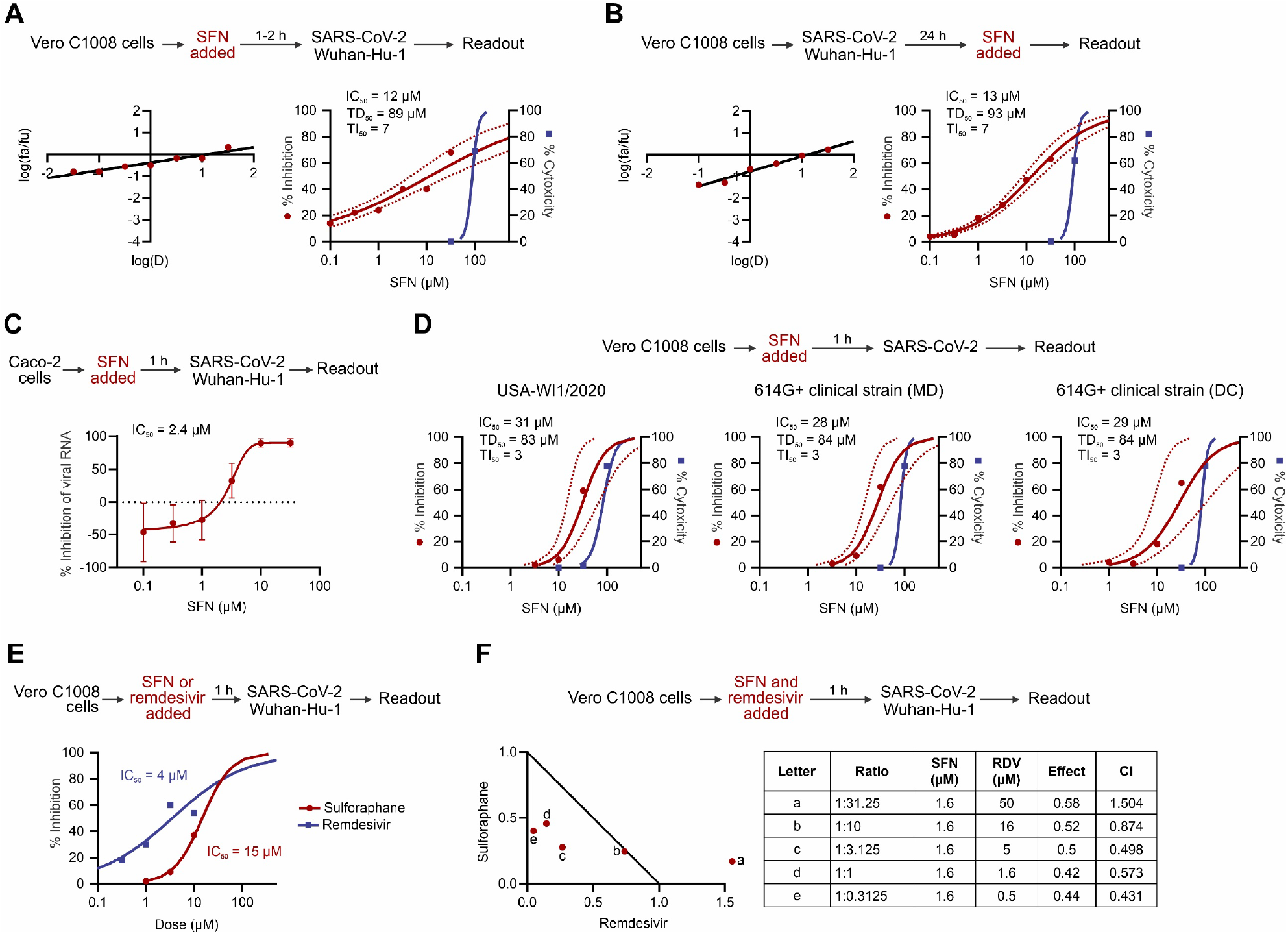
Antiviral effects of SFN against SARS-CoV-2. Median effect plot and dose-effect curves calculated for (**A**) Vero C1008 cells infected with SARS-CoV-2/Wuhan-Hu-1 after 1-3h incubation with increasing concentrations of SFN; (**B**) Vero C1008 cells infected with SARS-CoV-2/Wuhan-Hu-1 for 24h and then incubated with SFN. Antiviral data displayed in red; anti-host cell activity (cytotoxicity) displayed in blue. (**C**) The antiviral activity in human Caco-2 cells was determined by measuring viral RNA by qPCR. The cells were incubated with SFN for 1h before viral inoculation. (**D**) Effects of SFN evaluated in Vero C1008 cells exposed to drug for 1h followed by viral inoculation. A reference strain (USA-WI1/2020) and two 614G+ clinical strains of SARS-CoV-2 were evaluated for cytopathic effects using a bioluminescence readout. (**E**) Effects of SFN and remdesivir evaluated in Vero C1008 cells exposed to drug for 1h followed by viral inoculation. (**F**) Normalized isobologram showing combination index (CI) for combinations of various doses of SFN and remdesivir. Synergism (CI < 1); Additive effect (CI=1); Antagonism (CI >1); SFN, Sulforaphane; RDV, Remdesivir. Dotted lines represent 95% confidence interval. Experiments performed a minimum of 2 times (range = 2 – 7), 3 – 6 replicates within each experiment, except experiment shown in **E** which was performed once.

We further evaluated SFN for activity against a second reference strain of SARS-CoV-2 as well as two clinical strains that carry the spike D614G (614G+) substitution that is found in the majority of variants of concern currently in circulation (Figure 2D) ^9^. SFN inhibited USA-WA1/2020, (IC_50_ = 31 μM, 95% CI 14.7 – 66.4) and the two 614G+ clinical strains, USA/MDHP-20/2020 (MD) and USA/DCHP-7/2020 (DC) (IC_50_ = 28 μM, 95% CI 14.9 – 52.9, and 29 μM, 95% CI 8.2 – 102.3, respectively), with comparable efficacy to that reported above for reference strain Wuhan-Hu-1.

We next investigated whether SFN could affect an established virus infection. As shown in Figures 1C and 2B, SFN effectively inhibited both an HCoV-OC43 and a SARS-CoV-2-Wuhan-Hu-1 infection that had been allowed to replicate for 24 hours before addition of drug. The IC_50_ for both viruses was in the lower micromolar range, 18 μM (95% CI 4 – 84.1) and 13 μM (95% CI 8.6 – 20), respectively. Interestingly, these results show that the anti-virus specific activity, i.e., the therapeutic index (TI), of SFN is similar whether the drug is added just before or 24 hours after virus inoculation (Figures 1A, 1C, 2A-B) suggesting an effect on both extracellular entry and intracellular post-entry viral processes. We also determined whether a single application of SFN could protect from the cytopathic effects (CPE) of subsequent viral infection lasting 4 days. As shown in Figure 1D, SFN pretreatment of Vero C1008 host cells resulted in measurable inhibition of HCoV-OC43 CPE with an IC_50_ = 21 μM (95% CI 9.3 – 49.3) and TI_50_ = 4.

Finally, we examined the potential synergistic effects of SFN combined with the anti-viral drug remdesivir, an inhibitor of viral RNA-dependent RNA polymerase recently reported to shorten the time to recovery in adults who were hospitalized with COVID-19 ^10^. As shown in Figure 2E, remdesivir effectively inhibits in vitro replication of SARS-CoV-2 (IC_50_ = 4 μM) as well as HCoV-OC43, albeit at a higher concentration (IC_50_ = 22 μM) (Figure 1E). In two-drug combination assays, SFN and remdesivir interacted synergistically at several combination ratios to inhibit replication of both HCoV-OC43 (Figure 1F) and SARS-CoV-2-Wuhan-Hu-1 (Figure 2F) at concentrations below the corresponding IC_50_ for each drug.

To evaluate the ability of SFN treatment to reduce viral titers and inflammation in vivo, K18-hACE2 transgenic male mice were inoculated intranasally with 8.4×10^5^ tissue culture infectious dose 50 (TCID_50_) of SARS-CoV-2/USA/WI1/2020 ^11^. SFN was administered daily via oral gavage to a subgroup of infected animals starting one day prior to viral inoculation (Figure 3A).

**Figure 3.**
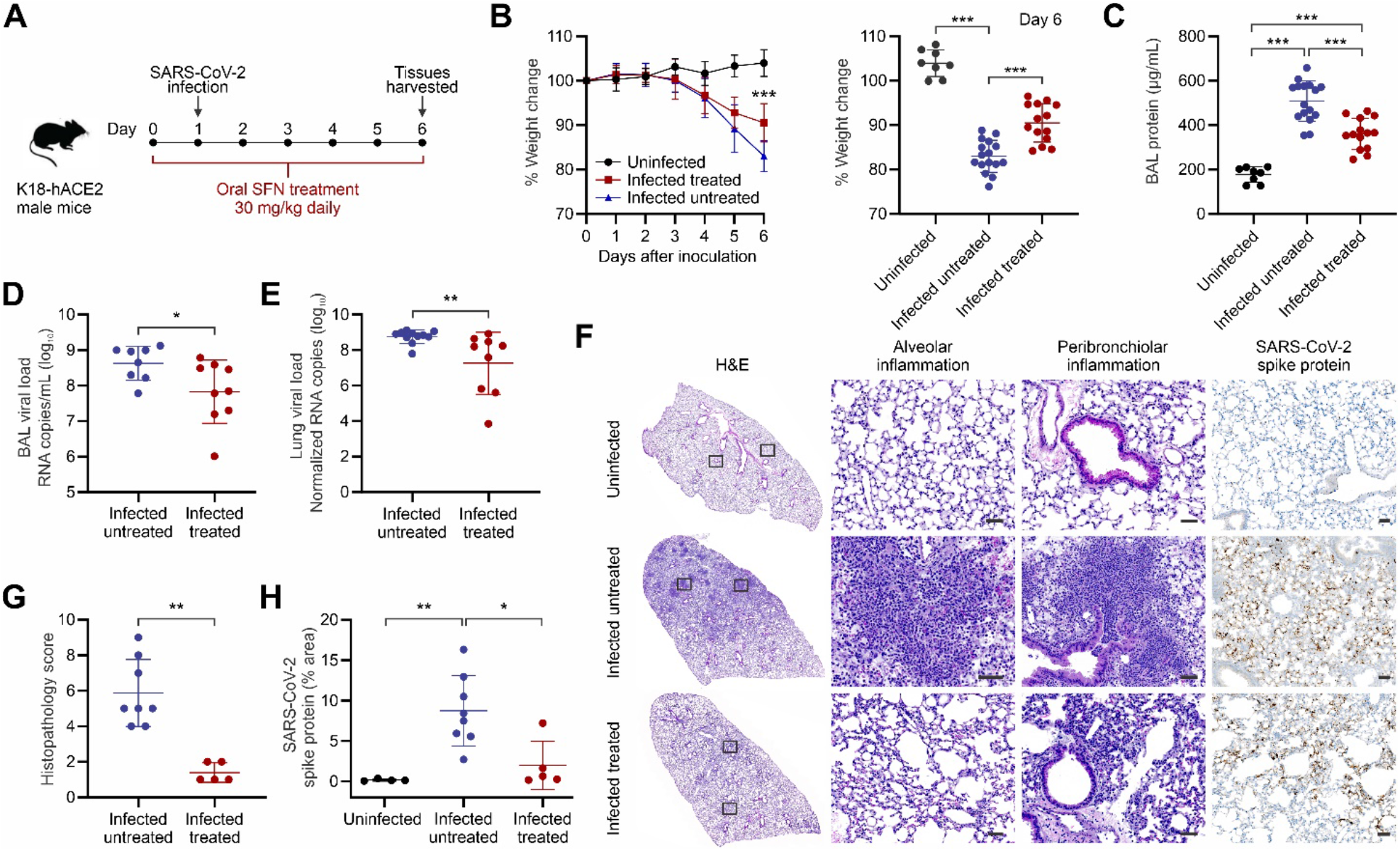
SFN treatment in SARS-CoV-2 infected K18-hACE2 mice. (**A**) Six-to eight-week-old male K18-hACE-transgenic mice were randomly distributed among treatment groups and inoculated intranasally with SARS-CoV-2/USA/WI1/2020 or vehicle. (**B**) Four days post inoculation, there was a marked weight loss in the infected groups although there was significantly less weight loss in the SFN treated animals. By day 6 post inoculation, the SFN-treated animals had lost 7.5% less bodyweight compared to infected untreated controls (one-way ANOVA, \*\*\**P*<0.0001).Data from three independent experiments, uninfected (n=8), infected untreated (n=16), infected treated (n=14). (**C**) Bronchoalveolar lavage (BAL) total protein quantification, determined as a surrogate for lung injury, measured 6 days post-infection. Infected untreated animals had significantly higher total protein compared to the infected treated group (one-way ANOVA, \*\*\**P*<0.0001). Data from three independent experiments, uninfected (n=8), infected untreated (n=16), infected treated (n=14). (**D**) The viral load in the BAL, as determined by qPCR, was significantly higher in infected untreated animals compared to the infected treated group (Mann-Whitney *U* test, two-tailed, \**P*=0.04). Data from two independent experiments, infected untreated (n=8), infected treated (n=9). (**E**) The viral load in the lungs of infected treated animals, represented as the SARS-CoV-2 N protein copies normalized to *Pol2Ra*, had a 1.5 log reduction compared to infected untreated controls (Mann-Whitney *U* test, two-tailed, \*\**P*=0.004). Data from two independent experiments, infected untreated (n=11), infected treated (n=9). (**F**) Hematoxylin and eosin (H&E) staining and immunostaining for SARS-CoV-2 spike protein, of histological sections of the lungs of a representative uninfected control, an infected untreated, and an infected treated mouse. Regions of the lung anatomy where alveolar and peribronchiolar inflammation was assessed are highlighted in boxes. Images show low (left panels; scale bar, 1mm) and high-power magnification (right panels; scale bar, 50μm) of the same tissue section. (**G**) Histopathological severity scoring was evaluated according to the pathological changes outlined in the methods section. Data from one independent experiment, infected untreated (n=8), infected treated (n=5). Mann-Whitney *U* test, two-tailed, \*\**P*=0.0008. (**H**) Quantification of the SARS-CoV-2 spike protein immunostaining showed a 4.41x lower % area in the lungs of SFN-treated mice compared to infected untreated controls (*P*=0.01). Data from one independent experiment, uninfected (n=4), infected untreated (n=8), infected treated (n=5). One-way ANOVA, \**P*<0.05, \*\**P*<0.005. All the data in this figure are represented as mean ± standard deviation, each dot represents one animal.

A marked weight loss was observed in the infected animals starting at four days post inoculation. By day 6 post inoculation, SFN-treated mice lost significantly less weight compared to controls (Figure 3B, *P*<0.0001). As a measure of lung injury, the protein concentration in the bronchoalveolar lavage (BAL) was significantly lower in the SFN-treated infected mice compared to untreated infected controls (Figure 3C, *P*<0.0001) suggesting a measure of protective effect of drug pretreatment. The viral burden measured in the alveolar fluid was also significantly lower in treated animals compared to untreated controls, with a 1.15 log reduction in viral titers (Figure 3D, *P*=0.04). Similarly, a 1.5 log reduction in viral lung titers was observed in SFN-treated mice compared to untreated controls, when normalized to *Pol2Ra* (Figure 3E, *P*=0.004). Data on pulmonary viral burden without normalization are presented in figure S1. Analysis of hematoxylin and eosin-stained lung sections from these animals showed an inflammatory process similar to what has been previously described for this model after SARS-CoV-2 infection ^11–13^ (Figure 3F). SFN-treated mice had a lower degree of pulmonary pathology with less alveolar and peribronchiolar inflammation, compared to infected untreated mice (Figure S2). Histopathology analysis showed a significant reduction a lung inflammation in SFN-treated mice (histopathology score of 1/16) over untreated controls (histopathology score of 6/16) (Figure 3G, *P*=0.0008). Immunostaining for SARS-CoV-2 spike protein revealed a more widespread distribution in the lungs of infected untreated animals compared to a focal distribution in those of treated animals (Figures 3F, S2). Quantification of the SARS-CoV-2 spike protein immunostaining revealed that the lung area associated with the virus was 4.4x higher in the infected untreated animals compared to the SFN-treated mice (Figure 3H, *P*=0.01).

Given the known immunomodulatory effects of SFN, we employed high-dimensional flow cytometry to evaluate the changes in the immune response of SARS-CoV-2-infected mice treated with SFN and untreated controls, as compared to uninfected mice (Figure S3, Table S1). Although the immunological landscape was altered as a result of the infection, there were limited differences in overall immune cell composition in the spleen or lungs between treated and untreated mice as visualized using Uniform Manifold Approximation and Projection (UMAP) (Figure 4A). While changes in the systemic immune responses reflected in the spleen were minimally different (Figure 4B-C), there were more pronounced effects locally within the lung (Figure 4B,D). Notably, infection induced significant recruitment of myeloid cells including monocytes and dendritic cells into the lungs of infected mice, however SFN-treatment significantly reduced this recruitment compared to infected untreated mice (Figure 4D, *P*<0.04). Recruitment of blood monocytes into the lung is known to initiate and maintain lung inflammatory responses, including ARDS, and has been demonstrated in SARS-CoV-2 infection ^14^. SFN treatment significantly decreased the percentage monocytes and CD11c+ dendritic cells out of total CD45+ immune cells in the lungs (Figure 5A, *P*=0.01). Further, metabolically distinct CPT1a+VDAC+ myeloid cells that have been shown to correlate with disease severity in patients with COVID-19 ^15^ were significantly decreased in response to SFN treatment (Figure 5A, *P*=0.01). Alveolar and interstitial macrophages from the lung of SFN-treated mice displayed lower expression of activation markers such as CD80, CD86, PD-L1 and MHC-II (Figure 5B, S4A-B). Activation also induced significantly lower frequencies of lung alveolar and interstitial macrophages producing cytokines such as IL-10, IL-1β, TNF-α, and TGF-β (Figure 5C, S4B). These findings were largely replicated in the bronchoalveolar lavage (Figure S5). Together, these data highlight the overall reduction of the local myeloid immune responses within the lung microenvironment as a result of SFN treatment. In line with the myeloid compartment, T cell activation was also diminished in response to SFN treatment. Directly ex vivo, CD8+ and CD4+ T cells isolated from the lung of infected untreated mice demonstrated increased expression of activation markers PD1 and MHC-II and the proliferation marker Ki-67, all of which were significantly decreased in SFN-treated mice (Figure 5D). This effect on T cell activation was predominantly seen in the lungs and was not found systemically in the spleen. Following stimulation with PMA/ionomycin, CD4+ T cells from the lung, but not the spleen, produced lower levels of IFN-γ and IL-10, however, the frequency of IL-4 and IL-17 were not significantly altered (Figure 5E, S6A-B). In summary, the immune-modulatory effects of SFN had a local effect of limiting immune cell activation within the lung, without disturbing or significantly altering systemic immune responses in the spleen.

**Figure 4.**
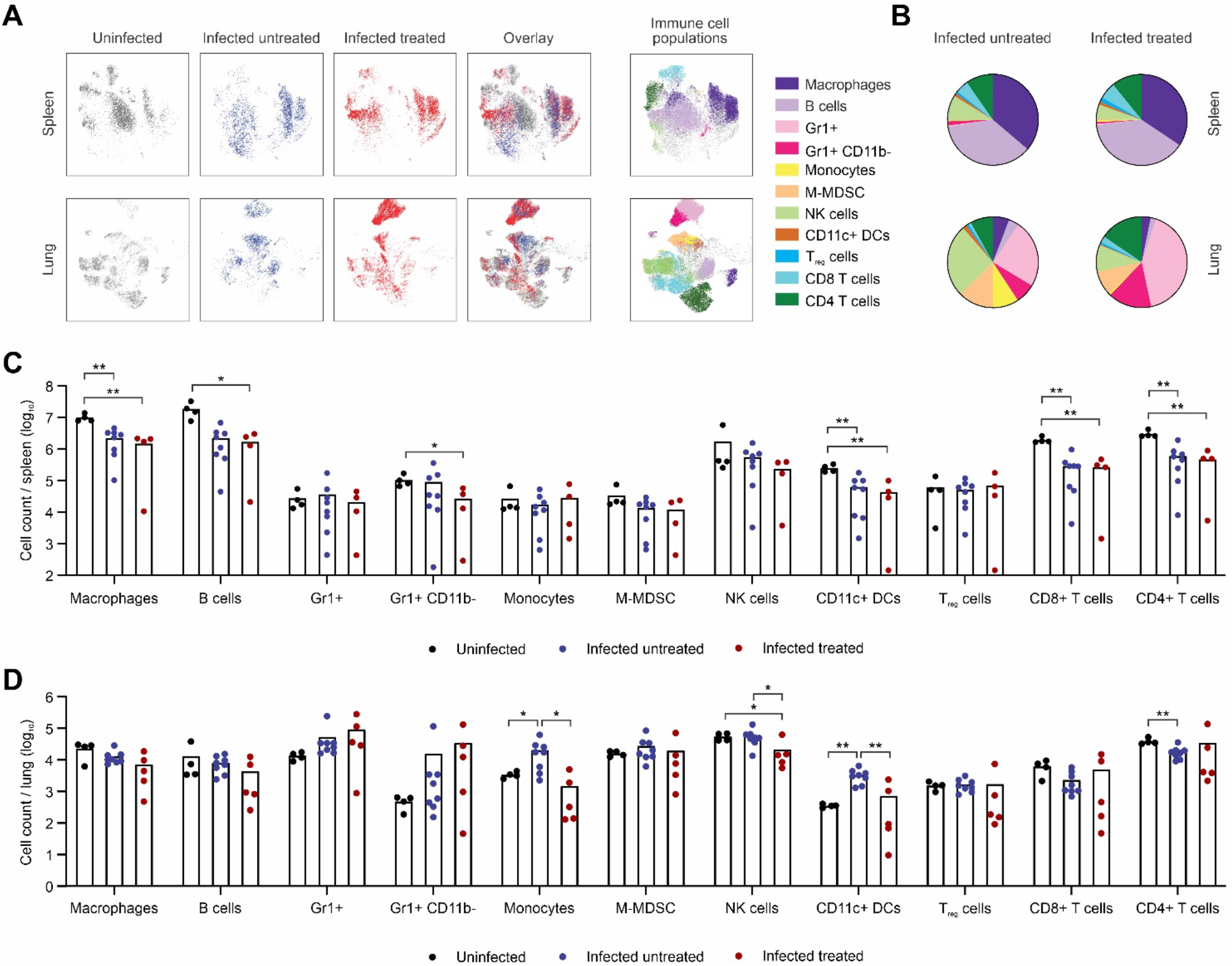
Effects of SFN treatment in the immune response. (**A**) Uniform Manifold Approximation and Projection (UMAP) was used to visualize all of the immune cell populations within the spleen and lung of uninfected (grey), infected untreated (blue), and infected treated (red) mice. The corresponding immune cell populations are presented in multiple colors in the panels on the right (**B**) Summary of immune cell frequencies out of total CD45+ immune cells in spleen and lung of infected treated or untreated mice. (**C**) Total cell count of indicated immune cell subset per spleen. Each dot represents one mouse. (**D**) Total cell count of indicated immune cell subset per lung. Each dot represents one mouse, data from one independent experiment. Bars represent mean values. DCs, dendritic cells; NK, natural killer, M-MDSC, mononuclear myeloid-derived suppressor cells. Statistical comparisons made with two-way ANOVA, \**P*<0.05, \*\**P* < 0.01.

**Figure 5.**
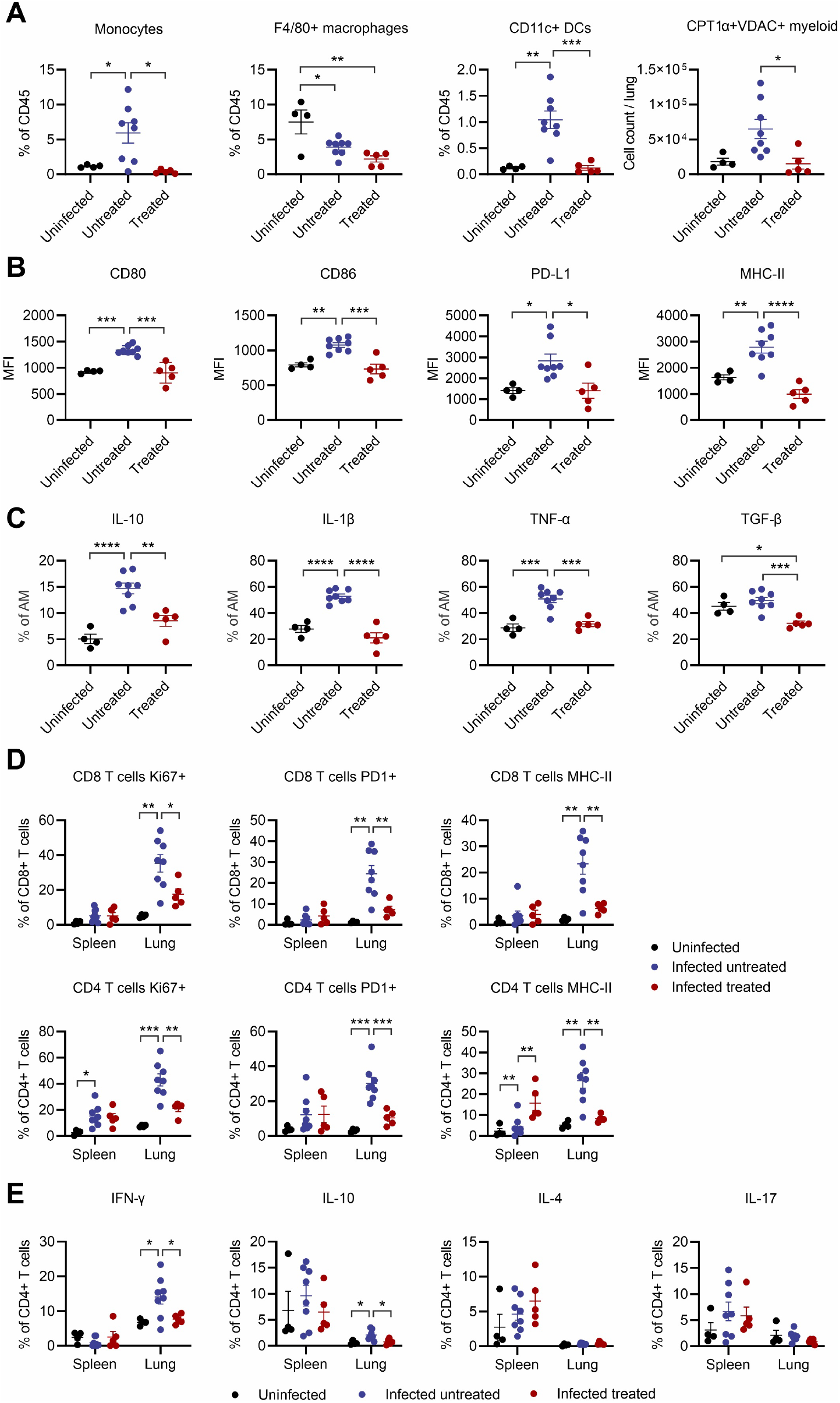
Functional characterization of the immune response after SFN treatment. (**A**) Myeloid cell subsets shown as percent of total CD45+ immune cells within the lung. (**B**) Alveolar macrophages (AM) after stimulation with protein transport inhibitors. MFI, mean fluorescent intensity. (**C**) Cytokine expression in alveolar macrophages after stimulation with protein transport inhibitors. (**D**) T cells were stained immediately ex vivo without further stimulation and evaluated for the expression of Ki67, PD1 and MHC-II. Percent of CD8+ or CD4+ T cells from spleen or lung expressing indicated marker are shown. (E) Percent of CD8+ or CD4+ T cells from spleen or lung expressing indicated marker after stimulation with PMA/ionomycin. Each dot represents one mouse, data from one independent experiment. Data represented as mean ± standard error of mean. n=4 uninfected, n=5 infected SFN-treated, and n=8 infected untreated animals. Statistical comparisons made with one-way ANOVA, \**P*<0.05, \*\**P* < 0.01, \*\*\**P* < 0.001, \*\*\*\**P* < 0.0001.

## Discussion

The ongoing SARS-CoV-2 pandemic has created the immediate need for effective therapeutics that can be rapidly translated to clinical use. Despite the introduction of vaccines, effective antiviral agents are still necessary, particularly considering the potential effects of viral variants^16^. While great efforts have been made to develop drugs that target the virus, this approach can be affected by the emergence of viral variants that change the affinity of the drug to the viral protein ^17^. An alternative approach is to also target host mechanisms required by the virus to infect cells and replicate ^18^. Host-directed therapy is advantageous as it allows pre-existing drugs to be repurposed, may provide broad-spectrum inhibition against multiple viruses, and is generally thought to be more refractory to viral escape mutations ^19,20^.

Following a preliminary examination of a small group of clinically approved drugs, experimental novel compounds, and nutritional supplements, SFN was identified as a promising candidate to target the host cellular response, given that it is orally bioavailable, commercially available, and has limited side-effects ^8,21^. We observed that SFN has dual antiviral and anti-inflammatory properties against coronaviruses. We determined that SFN has potent antiviral activity against HCoV-OC43 and multiple strains of SARS-CoV-2, with limited toxicity in cell culture. The similar results observed between the coronaviruses evaluated suggest that SFN could have broad activity against coronaviruses, a feature that may prove invaluable as new strains of pathogenic coronaviruses enter the human population. Moreover, synergistic antiviral activity was observed in vitro between SFN and remdesivir against both types of coronaviruses tested; comparable synergism in vivo would be advantageous in clinical scenarios where remdesivir is currently being used. We demonstrated in vivo efficacy of prophylactic SFN treatment using the K18-hACE2 mouse model of SARS-CoV-2 infection ^12^. Prophylactic SFN-treatment in animals reduced viral replication in the lungs by 1.5 orders of magnitude, similar to that reported for remdesivir in the same mouse model ^22^. As expected, SFN treatment also modulated the inflammatory response in SARS-CoV-2-infected mice, leading to decreased lung injury.

The pathogenesis of many viral infections is associated with increased production of reactive oxygen species (ROS) which leads to cell death ^23,24^. Conversely, SFN increases antioxidant, anti-inflammatory, and antiviral defenses primarily via activation of the cap‘n’collar transcription factor NRF2 ^25^. Under normal conditions, NRF2 remains in an inactive state by association with its inhibitor protein Kelch-like ECH-associated protein 1 (KEAP1) ^26^. In response to oxidative stress, KEAP1 is inactivated and NRF2 is released to induce NRF2-responsive genes that subsequently protect against stress-induced cell death ^27^. SFN has been extensively studied in humans for its anti-cancer properties, has been shown to activate the NRF2 pathway in upper airways ^28^, and improves the phagocytic ability of alveolar macrophages ^3^. The dual antiviral and anti-inflammatory properties of SFN have also been previously described for other viral infections. In vitro antiviral activity has been reported against influenza virus ^29^ and SFN treatment significantly limited lung viral replication and virus-induced inflammation in respiratory syncytial virus-infected mice ^30^.

Targeting the NRF2 pathway has been considered a promising approach to develop therapeutics for COVID-19 for multiple reasons ^31^. NRF2 deficiency is known to upregulate the angiotensin-converting enzyme 2 (ACE2), the primary mechanism of cell entry for SARS-CoV-2. The NRF2 activator oltipraz reduces ACE2 levels, suggesting that NRF2 activation might reduce the availability of ACE2 for SARS-CoV-2 entry into the cell ^32^. Increased NRF2 activity also reportedly inhibits IL-6 and IL-1β gene expression ^33^, two cytokines known to play key roles in promoting the hyperactive immune response in severely ill COVID-19 patients ^34^. Conversely, NRF2 activity is dysregulated in disease states that have been associated with increased severity of COVID-19, e.g. diabetes ^35^. Further, NRF2 activity declines in older patients who are more susceptible to severe COVID-19 ^36^. Recent reports suggest that NRF2-dependent genes are suppressed in SARS-CoV-2 infected cells and in lung biopsies from COVID-19 patients ^31^. Similarly, treatment of cells with NRF2 agonists 4-octyl-itaconate and dimethyl fumarate inhibited replication of SARS-CoV-2 in vitro ^31^. SFN also inhibits inflammation through NRF-2 independent pathways, such as reducing the pro-inflammatory nuclear factor kappa B (NF-κB) ^37^. NF-κB activation has been described as a key component of the inflammatory response to multiple viral infections, including COVID-19 ^38^. There are also other pathways affected by SFN (e.g. STING) that could play a role in its antiviral response to coronaviruses. These additional pathways are the subject of continuing investigation ^39^.

As a potent NRF2 activator, SFN can modulate the host’s immune response while also providing direct antiviral effects. In contrast to therapeutics that inhibit a single cytokine (e.g. IL-6, IL-1β, etc.) ^40^, SFN has important and diverse effects in modulating the lung immune response to SARS-CoV-2 infection. Excessive inflammatory response to SARS-CoV-2 leads to severe disease or death in patients with COVID-19 ^41^. Therefore, promoting a balanced and robust antiviral response while modulating excessive innate inflammatory responses could represent a favorable scenario that could reduce viral load while also limiting collateral damage to the infected lung. As has been previously reported, SARS-CoV-2 infection leads to an increase in pulmonary dendritic cells and a reduction in CD4+ T cells in K18-hACE2 mice ^12^. We also observed substantial accumulation of immune cells in the lungs of SARS-CoV-2 infected mice, consistent with what has been noted on postmortem analysis of patients with COVID-19 ^12,42^, as well as decreased numbers of T cells in the spleen, consistent with human studies where lymphopenia is correlated with severe COVID-19 ^43^. SFN treatment had significant effects on multiple immune cell populations in the lungs, with a reduction in monocytes, NK cells, and dendritic cells compared to infected untreated controls. These findings are likely the effect of a combination of the overall reduced inflammation and direct effects of SFN on specific cell populations. For example, NK cells exposed to SFN had increased cell lytic function through dendritic cell-mediated IL-12 production ^44^. We found decreased recruitment of myeloid cells to the lungs of treated mice and decreased activation profile of local macrophages. The presence of alveolar macrophages with transcriptionally upregulated inflammatory genes and increased secretion of IL-1β have been associated with worse outcomes and increased mortality in patients with ARDS ^14,45–47^. Our results show increased IL-1β in alveolar macrophages with SARS-CoV-2 infection, which was abrogated by SFN treatment (*P*<0.0001). Mechanistically, the benefits of SFN therapy in our model could be due in part to its modulatory effects on myeloid cells after SARS-CoV-2 infection. SFN treatment led to a reduction in TNF-α in alveolar macrophages and IFN-γ in T cells, both of which are key triggers of cell death and mortality in SARS-CoV-2 infection and cytokine shock syndromes ^48^. Further, SFN was able to reduce, but not eliminate T cell activation within the lung. This reduction in T cell activation could be a direct effect in T cells or could operate through downregulation of myeloid costimulatory for T cells such as CD80/CD86. SFN might therefore be able to modulate and dampen immune responses without inhibiting immunity necessary for viral clearance.

While the K18-hACE2 mouse model has been previously used to recapitulate features of COVID-19 in humans ^12^, our study has several limitations. The expression of the hACE2 transgene is non-physiological, resulting in tissue expression levels that are distinct from endogenously expressed ACE2. Sex differences, which are known to occur with SARS-CoV-2 infection, could not be assessed since we only used male animals in these experiments. Finally, the absorption of SFN after oral administration can be modified by the intestinal microbiome ^1^, leading to potentially variable drug exposures between animals.

Our results demonstrate that pharmacologically relevant micromolar concentrations of SFN inhibited viral replication and virus-induced cell death. The bioavailability of SFN in humans is dependent on dosage and dietary form ^1^. Safety studies using SFN-rich broccoli sprouts corresponding to 50 to 400 μM SFN daily have shown that SFN is well tolerated without clinically significant adverse effects ^1,21,49,50^. After intake, the plasma concentration of SFN is rapidly eliminated but SFN exerts a sustained effect on gene expression ^51^. Consumption of SFN-rich broccoli sprouts (single oral daily dose equivalent to 200μM of SFN) reportedly results in a peak plasma concentration (C_max_) of 1.9 μM at 2-3h ^52,53^. By comparison, SFN inhibited in vitro SARS-CoV-2 replication in human cells with an IC_50_ of 2.4μM. Doses of 100μM of SFN administered twice daily led to higher steady-state concentrations of SFN in plasma, compared to a single dose ^1,52,54^. In summary, our results suggest that SFN could be a rapidly applicable strategy for the prevention and treatment of COVID-19, as well as seasonal coronavirus infections.

## Conclusion

We documented that SFN can inhibit the in vitro and in vivo replication of SARS-CoV-2 at pharmacologically and therapeutically achievable doses and can modulate the inflammatory response thereby decreasing the consequences of infection in the animal model when administered prior to infection. Given that SFN is orally bioavailable, commercially available, and has limited side-effects, our results suggest it could be a promising and easily scalable approach for the prevention and treatment of COVID-19 as well as other coronavirus infections.

## Materials and Methods

### Drugs

L-SFN, 10mg/mL in ethanol (56mM), was obtained from Cayman Chemical (Ann Arbor, MI). D,L-SFN was obtained from Millipore Sigma (St. Louis, MO); stock solution of 5mM was prepared in DMSO. Remdesivir was obtained from MedChemExpress or Cayman Chemical and stock solutions, 5 or 20mM, respectively, were prepared in DMSO. Drug stock solutions were stored at −25°C.

### Cells and viruses

All cells were obtained from the American Type Culture Collection (ATCC, Manassas, VA, USA). HCT-8 [HRT-18] (ATCC CCL-244) and Vero C1008 [Vero 76, clone E6, Vero E6] (ATCC CRL-1586) cells were used for growing virus and determining virus stock titers. Vero C1008 cells, MRC-5 (ATCC CCL-171) cells, and Caco-2 (ATCC HTB-37) cells were used as host cells in antiviral assays. HCT-8 cells were grown in RPMI-1640 medium supplemented with 10% fetal bovine serum (FBS) (MilliporeSigma, St. Louis, MO, USA), L-glutamine, penicillin-streptomycin, and sodium pyruvate. Vero C1008 and MRC-5 cells were grown in EMEM with 10% FBS, L-glutamine, and penicillin-streptomycin at 37°C with 5% CO_2_. Caco-2 cells were grown in Minimum Essential Media supplemented with 10% FBS, 1x sodium pyruvate and penicillin-streptomycin at 37°C with 5% CO_2_. Human coronavirus OC43 (HCoV-OC43) was purchased from ATCC (Betacoronavirus 1, ATCC VR-1558). SARS-CoV-2/Wuhan-1/2020 virus (U.S. Centers for Disease Control and Prevention) was provided to us by Dr. Andrew Pekosz. 2019-nCoV/USA-WA1/2020 was obtained through BEI Resources, National Institute of Allergy and Infectious Diseases (NIAID), National Institutes of Health (NIH). Two 614G+ clinical strains of SARS-CoV-2, SARS-CoV-2/USA/DCHP-7/2020 (DC), and SARS-CoV-2/USA/MDHP-20/2020 (MD) were isolated from patients at The Johns Hopkins Hospitals ^55^. The virus stocks were stored at −80°C and titers were determined by tissue culture infectious dose 50 (TCID_50_) assay. All work with infectious SARS-CoV-2 was performed in Institutional Biosafety Committee approved BSL3 and ABSL3 facilities at Johns Hopkins University School of Medicine using appropriate positive pressure air respirators and protective equipment.

### CPE inhibition assay

We used a colorimetric assay that interrogates both antiviral and anti-host cell activities to evaluate compounds ^56^. This assay is predicated upon the virus’s ability to cause a cytopathic effect (CPE), measured in TCID_50_. Host cells, 7.5-10 × 10^3^ in virus growth medium (VGM; Dulbecco’s Modified Eagle Medium without phenol red supplemented with 3% FBS), were plated in clear 96-well half-area tissue cultures plates or white, clear-bottom 96 well plates, 24 hours prior to the assay. On the day of the assay, working solutions of drugs (0.1 – 1mM) were made by dilution of drug stocks in VGM. For one-drug analysis assays, 50 μL of the drug working solution was added to each well in the first column of cells and then drugs were serially diluted across the plate by dilutions of 0.5 log_10_. The default drug test range was 320 – 0.032 μM. Drug-exposed cells were incubated for 1 – 24 hours at 35°C (HCoV-OC43) – 37°C (SARS-CoV-2), after which time 32-50 TCID_50_ of virus suspended in VGM or VGM alone was added to cells. Test plates had virus control wells (virus + / drug −), drug control wells (virus − / drug +), and cell control wells (virus − / drug −). After 3 – 4 days incubation at 35°C – 37°C / 5% CO_2_, the cell viability was assessed using Celltiter 96®AQ_ueous_ One Solution (Promega Corp, WI, USA) or CellTiter-Glo® One Solution Assay system (Promega) following manufacturer protocols. Color reactions were read at 490-650nm absorbance in a Filtermax F5 microplate reader (Molecular Devices, CA, USA) using SoftMax Pro 6.5 software. Luminescence readouts were obtained in a FLUOstar Omega plate reader (BMG Labtech, Ortenberg, Germany). For two-drug combination assays, one test drug was serially diluted across the plate (left to right) as described above; the second drug was serially diluted down the plate (top to bottom). The starting concentration of the first drug was adjusted to allow for dilution with the second drug. For interrogation of a drug’s ability to affect an established viral infection, host cells were infected with virus and allowed to incubate 24 hours. After this time, the VGM in the wells was removed, the cells rinsed with Dulbecco’s phosphate buffered saline, and then drug dilutions in the ranges mentioned above, or VGM only, were added to appropriate wells. Cell viability reagent was added at 3-4 days post-infection. For examination of a drug’s ability to prevent an infection by pretreatment of the host cells, the test drug was serially diluted across the plate as described above. After 24 hours incubation, the drug was removed by aspiration, the cells were rinsed once with warm Hanks Balanced Salt Solution, and then 32 TCID_50_ of virus in drug-free VGM was added to appropriate wells. Cell viability reagent was added at 4 days post-infection.

### In vitro data analysis

Calcusyn software (Biosoft, Cambridge, UK) was used to calculate the median inhibitory concentration (IC_50_), median cytotoxic dose (TD_50_), and to generate median effect plots and dose-response curves. The therapeutic index (TI), a measure of antiviral selectivity, was calculated by the formula TI = TD_50_ / IC_50_. Combination Indices (CI) for two-drug combination assays were calculated by Calcusyn software. Isobolograms to depict synergistic, additive, and antagonistic combinations were generated by the software.

### Viral RNA determination

Zymo Quick-RNA Viral 96 Kit (Zymo Research) was used to isolate RNA from cell supernatants according to the manufacturer’s protocol. cDNA synthesis was performed using qScript cDNA Supermix containing random hexamers and oligo-dT primers following the manufacture’s protocol (Quanta Biosciences). Real-time quantitative reverse transcription PCR (RT-qPCR) was performed in technical triplicate for each sample using TaqMan Fast Advanced Master Mix (Applied Biosystems) on a StepOne Plus Real Time PCR machine (Applied Biosystems). Primers and probes are listed below. The cycling parameters were as follows: (i) 2 min at 50 °C; (ii) 2 min at 95 °C; and (iii) 45 cycles at 95 °C for 3 s and 55 °C for 30 s. Molecular standard curves were generated using serial dilutions of a plasmid containing the complete SARS-CoV-2 N gene (Integrated DNA Technologies, Catalog #10006625). SARS-CoV-2 RNA was detected using premixed forward (5’-TTACAAACATTGGCCGCAAA-3’) and reverse (5’-GCGCGACATTCCGAAGAA-3’) primers and probe (5’-FAM-ACAATTTGCCCCCAGCGCTTCAG-BHQ1-3’) designed by the U.S. CDC as part of the COVID-19 CDC Research Use Only kit (Integrated DNA Technologies, Catalog #: 10006713) to amplify a region of the SARS-CoV-2 nucleocapsid (N) gene. For lung tissues, the sample was homogenized in 1mL of TRIzol (Invitrogen) and the RNA was isolated using a combined protocol of TRIzol phenol chloroform and the RNeasy Mini Kit (Qiagen) according to the manufacturer’s protocol, and RT-qPCR was performed as described above. For lung tissue lysates, viral copies per lung sample were normalized to the relative expression of the mouse RNA Polymerase II gene (*Pol2Ra*) using the TaqMan gene expression assay (Catalog #: Mm00839502_m1; ThermoFisher) ^57^.

### Animal experiments

Animal studies were carried out based on the recommendations in the Guide for the Care and Use of Laboratory Animals of the National Institutes of Health. The protocols were approved by the Johns Hopkins University Institutional Animal Care and Use Committee. Heterozygous K18-hACE2 C57BL/6J mice (strain: 2B6.Cg-Tg(K18-ACE2)2Prlmn/J) were obtained from The Jackson Laboratory and propagated at Johns Hopkins University School of Medicine. Animals were separately housed in groups and fed standard chow diets. Male mice, 6-8 weeks old were used for this study. A subgroup of animals received 30mg/kg daily of SFN diluted in 2% ethanol in water via oral gavage. Treatment started one day prior to viral infection. Infected untreated and uninfected controls also received daily oral gavage with 2% ethanol in water. After induction of anesthesia with ketamine hydrochloride and xylazine, the animals received 8.4×10^5^ TCID_50_ of SARS-CoV-2/USA/WI1/2020 intranasally. Uninfected animals received intranasally the same volume of vehicle (2% ethanol and water). Weights were monitored daily, the animals were sacrificed 6 days post-infection by isoflurane overdose, and the tissues were harvested. Tissues were perfused with PBS after serum collection via cardiac puncture and before tissue harvest. Broncheoalveolar lavage (BAL) was obtained by cannulating the trachea with a 20-gauge catheter. The right lung was lavaged twice (each aliquot 1 ml; calcium-free PBS); total returns averaged 1–1.5 ml/mouse. BAL was centrifuged at 600 g for 8 minutes at 4°C. The cell-free supernatants were stored at –80°C for total protein quantification using the BCA protein assay (Sigma).

### Flow cytometry

Lungs were minced and incubated at 37°C in an enzyme cocktail of RPMI containing 2.4 mg/ml collagenase I and 20 μg/ml DNase (Invitrogen), then mashed through a 70-μm nylon cell strainer (BD Falcon). All flow cytometry antibodies used for phenotypic and metabolic analysis can be found in Table S1. For analysis immediately ex vivo, cells were washed once in PBS and immediately stained for viability with Biolegend Live/Dead Zombie NIR Fixable Viability Dye and Fc Block for 10 min at room temperature. Cell surface staining was performed in 100μL of 20% BD Horizon™ Brilliant Stain Buffer + PBS with surface stain antibody cocktail for 20 min at room temperature. Cells were fixed and permeabilized with eBioscience™ FoxP3/Transcription Factor Staining kit 1x Fixation/Permeabilization reagent overnight at 4°C. Cells were washed with 1x Permeabilization/Wash buffer. Intracellular staining (ICS) was performed in 100μL 1x Permeabilization/Wash buffer with ICS antibody cocktail for 45 min at room temperature. Cells were washed once with Permeabilization/Wash buffer then resuspended in Permeabilization/Wash buffer for acquisition by flow. To improve the quality of the T-cell flow cytometry functional staining, the cells were stimulated with phorbol 12-myristate 13-acetate (PMA, 50ng/mL) and inomycin (1μg/mL) for 1h, following for a 3h incubation with protein transport inhibitors (GolgiPlug and GolgiStop, BD). For the myeloid flow cytometry functional staining, the cells were incubated only with protein transport inhibitors for 4h. Samples were run on a 3 laser Cytek Aurora spectral flow cytometer or a FACSAria II spectral flow cytometer (BD). FCS files were analyzed using Flowjo v10.6.2 software (BD). Manual gating strategies for all the panels can be found in Figure S3. High-dimensional unbiased analysis of cell phenotypes was performed using Flowjo plugins DownSample v3 and UMAP.

### Histology and immunohistochemistry

After euthanasia, tissues were fixed on 10% neutral-buffered formalin. Tissues were embedded in paraffin and sections were stained with hematoxylin and eosin. Digital light microscopy scans of the lung were examined implementing a semi-quantitative, 5-point grading scheme (0 - within normal limits, 1 - mild, 2 - moderate, 3 - marked, 4 -severe) based on what has been previously reported by White *et al*. for this infection model ^22^. The scoring system considered four different histopathological parameters: 1) perivascular inflammation 2) bronchial or bronchiolar epithelial degeneration or necrosis 3) bronchial or bronchiolar inflammation and 4) alveolar inflammation. These changes were absent (grade 0) in lungs from uninfected mice. Individual total pathology scores varied from 1/16 to 2/16 for the SFN-treated group, and from 4/16 to 9/16 for the infected untreated controls. Immunostaining was performed at the Oncology Tissue Services Core of Johns Hopkins University School of Medicine. Immunolabeling for the SARS-CoV-2 spike protein was performed on formalin-fixed, paraffin-embedded sections on a Ventana Discovery Ultra autostainer (Roche Diagnostics). Briefly, following dewaxing and rehydration on board, epitope retrieval was performed using Ventana Ultra CC1 buffer (catalog # 6414575001, Roche Diagnostics) at 96°C for 64 minutes. Primary antibody, anti-SARS-CoV-2 spike protein (1:200 dilution; catalog # GTX135356, lot # 43957, Genetex) was applied at 36°C for 60 minutes. Primary antibodies were detected using an anti-rabbit HQ detection system (catalog # 7017936001 and 7017812001, Roche Diagnostics) followed by Chromomap DAB IHC detection kit (catalog # 5266645001, Roche Diagnostics), counterstaining with Mayer’s hematoxylin, dehydration, and mounting. Automated analysis of the SARS-CoV-2 spike protein immunostaining was performed with Halo v3.2.1851.328 (Indica Labs). The area corresponding to the anti-SARS-CoV-2 spike protein was divided by the sum of the areas corresponding to cellular structures stained with Mayer’s hematoxylin + anti-SARS-CoV-2 spike protein. The ratio is represented as a percentage.

### Statistical analysis

Data were analyzed using Prism 9.0.1 (GraphPad). Specifics of statistical comparisons are detailed in individual figure legends. Statistical significance was assigned when *P* values were <0.05.

## Acknowledgments

We thank Dr. Andrew S. Pekosz for providing the viral strains, and Sujayita Roy for the assistance with IHC staining. We also thank Dr. Jason Villano for supporting animal breeding and Drs. Jed W. Fahey and Craig W. Hendrix for valuable expert discussions about this project and manuscript (JWF).

## Funding

This study was supported by the National Institutes of Health (R01-AI153349, R01-AI145435-A1, and R21-AI149760 to S.K.J. and A.A.O.), support from the Center for Infection and Inflammation Imaging Research (Johns Hopkins University), Mercatus Center (Fast Grant #2167 to C.K.B.), and the Stanley Medical Research Institute (R.H.Y. and L.J-B.).

## Author contributions

Conceptualization: AAO, LJ-B. Formal Analysis: AAO, CKB, LJ-B, AFV-R, EAT, JDP, FRD. Funding acquisition: RHY, SKJ, LJ-B. Investigation: AAO, CKB, SLD, OK, AFV-R, MLT, EAT, LJ-B. Methodology: AAO, LJ-B. Resources: RHY, SKJ. Supervision: AAO, LJ-B, RHY, SKJ. Validation: AAO, LJ-B, RHY, SKJ. Visualization: AAO. Writing – original draft: AAO, LJ-B. Writing – review & editing: CKB, AFV-R, EAT, MLT, SLD, OK, JDP, FRD, RHY, SKJ.

## Competing interests

LJ-B, AAO, RHY, and SKJ are co-inventors on pending patent application USPA #63/142,598, “Methods for inhibiting coronaviruses using sulforaphane” filed by Johns Hopkins University.

## Data availability

All data associated with this study are present in the paper or the Supplementary Materials.

